# Effects of wasting disease on sea star populations in southern California: Variations over a 40-year period

**DOI:** 10.1101/2025.07.30.667779

**Authors:** John D. Dixon, Stephen C. Schroeter, Richard O. Smith, P.K. Dayton, P. Ed Parnell

## Abstract

Sea star wasting disease has been described for asteroid populations from many parts of the world and for numerous species. The causative agents are generally unknown. However, within the Southern California Bight, a *Vibrio* bacterium appears to have been the cause of wasting disease in the 1980s, and probably the 1990s, during warm water episodes associated with El Ninos. Although a densovirus was implicated in the mass mortality of one species of subtidal sea star during the geographically widespread epizootics that began in 2013, the etiology appears to vary among species, locations, and time periods, and in some cases may not even be associated with a pathogen.

We have been studying subtidal benthic communities in southern California since the 1970s and have documented the population effects of wasting disease epizootics associated with warm water El Niño events from 1980 through 2020. Until 2015, population declines coincident with wasting disease were followed by various degrees of recovery, whereas after 2015 there has been little or no recovery at our study sites. Prior to 2013, wasting disease in the northeast Pacific was only reported from locations in the Gulf of California, the southern California bight, and the coast of Vancouver Island, British Columbia. Since 2013, epizootics of wasting disease among sea stars have been observed along the entire west coast of North America. We speculate that the lack of recovery after 2015 may be due to a reduction in larval supply caused by the greater geographical extent of the disease.

## 1. INTRODUCTION

Wasting disease in echinoderms refers to a set of symptoms associated with a disease process characterized by necrosis that begins as lesions within the epidermis and soon spreads throughout the body, generally resulting in death. The disease can be remarkably virulent and fast-acting and can result in epizootics that cause mass mortality over large areas.

In the northeast Pacific, wasting disease in asteroids was first observed at numerous intertidal locations in the Gulf of California in June 1978 (Dungan, et al. 1982). At Puerto Peñasco at the head of the Gulf of California, the intertidal sun star, *Heliaster kubiniji*, suffered catastrophic mortality and, within two weeks, the population density plummeted from 0.1-1.0 m^-2^ to near zero. Everywhere observations were made in the Gulf of California, the sun star essentially disappeared, and after three years still occurred in low numbers in its rocky intertidal habitat. The only other species that was observed with wasting disease symptoms was the sea star *Othilia tenuispinus*. Where *O. tenuispinus* occurred in the intertidal zone it suffered heavy mortality. Subtidal individuals were not affected. This disease episode was correlated with record high seawater temperatures (Dungan et al. 1982).

During the summers of 1978 and 1979 dead and dying sea stars with necrotic lesions were also observed in intertidal and shallow subtidal habitats in southern California and on Catalina Island (Dungan, et al. 1982; personal observations and observations by J. Kastendiek, and J. Engel). Although the demographic effects were not documented, anecdotal observations suggest that the density of intertidal sea stars was significantly reduced. Two additional sea star epizootics occurred within the Southern California Bight during warm water episodes associated with the 1982-1983 and 1997-1998 El Niños. From 1981 to 1984, the density of sea stars in rocky subtidal habitats, in depths up to at least 18 m, declined markedly at our study sites in Southern California as a result of wasting disease. From 1997 to 1998, sea stars in the Channel Islands suffered a similar epizootic, with the most abundant species, *Asterina miniata*, suffering a 75% decline in density during the summer of 1997 (Eckert et al. 2000). Mass mortality was not apparent at this time at our mainland sites where sea star densities were still very low. Between epizootics, sea stars with the symptoms of wasting disease were occasionally observed both along the mainland coast and at the Channel Islands (personal observations; Eckert et al. 2000). Although animals with necrotic lesions were observed as far north as Point Estero, 16 km north of Morro Bay (Eckert et al. 2000), the three episodes of mass mortality along northeastern Pacific shores from wasting disease during the period 1978 to 1998 were mainly confined to the Southern California Bight and were associated with periods of unusually warm water. That the occurrence and progression of the disease that caused these epizootics was temperature-related is suggested by the observations that sea stars in deep water (> 30 m) were not affected (Tegner & Dayton 1987) and that wasting disease symptoms could be reversed by transferring diseased individuals to cold water (Eckert et al. 2000; personal observations).

A far more geographically widespread sea star epizootic began during a marine heat wave in 2013-2014. Although wasting disease had previously been observed as far north as Vancouver Island, British Columbia (Bates et al. 2009), mass mortality was first reported from Washington’s Olympic National Park in June 2013 (Schrope 2014). Wasting disease was soon documented among intertidal populations from Alaska to Baja California causing mortality in at least 13, and perhaps in as many as 22, asteroid species (https://www.eeb.ucsc.edu/pacificrockyintertidal/data-products/sea-star-wasting). Although subtidal populations were not as extensively monitored, those that were studied also suffered heavy mortality (Montecino-Latorre et al. 2016; Shultz et al. 2016; this study).

The episodes of sea star wasting disease observed in Washington and British Columbia appeared to be temperature related (Bates et al. 2009; Eisenlord et al. 2016). The 2013-2014 epizootic coincided with unusual warm seawater temperature anomalies (Bond et al. 2015). In laboratory experiments where *Pisaster ochraceous* was held at 19°, 16°, 14° and 12°C, the disease progressed more rapidly at the higher temperatures (Eisenlord et al. 2016). Similarly, *P. ochraceous* housed at 9°C lived twice as long as individuals held at 12°C (Kohl et al. 2016). However, to put these data in perspective, the average sea surface temperature in the San Juan Islands in August 2014 was 11.7°C (NOAA station 9449880, Friday Harbor) and in both laboratory experiments all animals held at 12°C died within 12 days, whereas the epizootics of the 1980s, and 1990s all occurred during the late summer and fall when sea surface temperatures exceeded 20°C, and sea stars with lesions did not die and began to heal when held in aquaria at 12°-13°C (personal observations).

Hewson et al. (2014), attempted to identify the agent that caused the wasting disease epizootic that began in 2013. They focused on viruses because they found no evidence of prokaryotic or eukaryotic microbial infection in seastars with wasting disease. Through challenge experiments with live and heat-killed inocula of virus-sized material, they demonstrated that the pathogen causing wasting disease in the sea star *Pycnopodia helianthoides* in their experiments was a virus-sized organism. Subsequent metagenomic analysis of tissue homogenates from *P. helianthoides* with and without disease symptoms from the challenge experiments demonstrated that the microorganism they named “sea star-associated-densovirus,” or SSaDV, was the sole virus associated with wasting disease in their samples. Based on a statistical analysis of the viral load and prevalence of SSaDV in field samples, they concluded that the densovirus was the most likely virus involved in the disease. However, in later work, that included challenge experiments with species other than *P. helianthoides*, there was no clear association between viruses and wasting disease in any species other than *P. helianthoides* and there was no consistent relationship with environmental variables such as sea water temperature (Hewson, et al. 2018).

The epizootic that began in 2013-2014 differed markedly from the earlier mass die-offs of sea stars from wasting disease. It occurred over a much larger area, it occurred at much lower temperatures in the north, and it originated in Washington and British Columbia rather than in southern waters. Another striking difference is that the epizootics of the 1980s, and possibly all the earlier epizootics in southern waters, were caused by a bacteria-sized microorganism, identified as a *Vibrio* species (this report), whereas the etiologies of the epizootics of 2013-2014 and later are generally unknown. However, it is unlikely that there is a single etiology or that it is the same disease among all the species and locations where it is observed (Hewson, et al. 2018). In some cases, wasting disease may not be caused by a pathogen but by depleted O_2_ conditions due to microbial activity (Aquino, et al., 2021.

## 2. MATERIALS AND METHODS

### 2.1 Sea Surface Temperature

Since 1916, sea surface temperatures have been recorded daily by Scripps Institution of Oceanography personnel at the SIO pier, about 18 km north of the Point Loma study sites (Figure 1).

**Figure 1.**
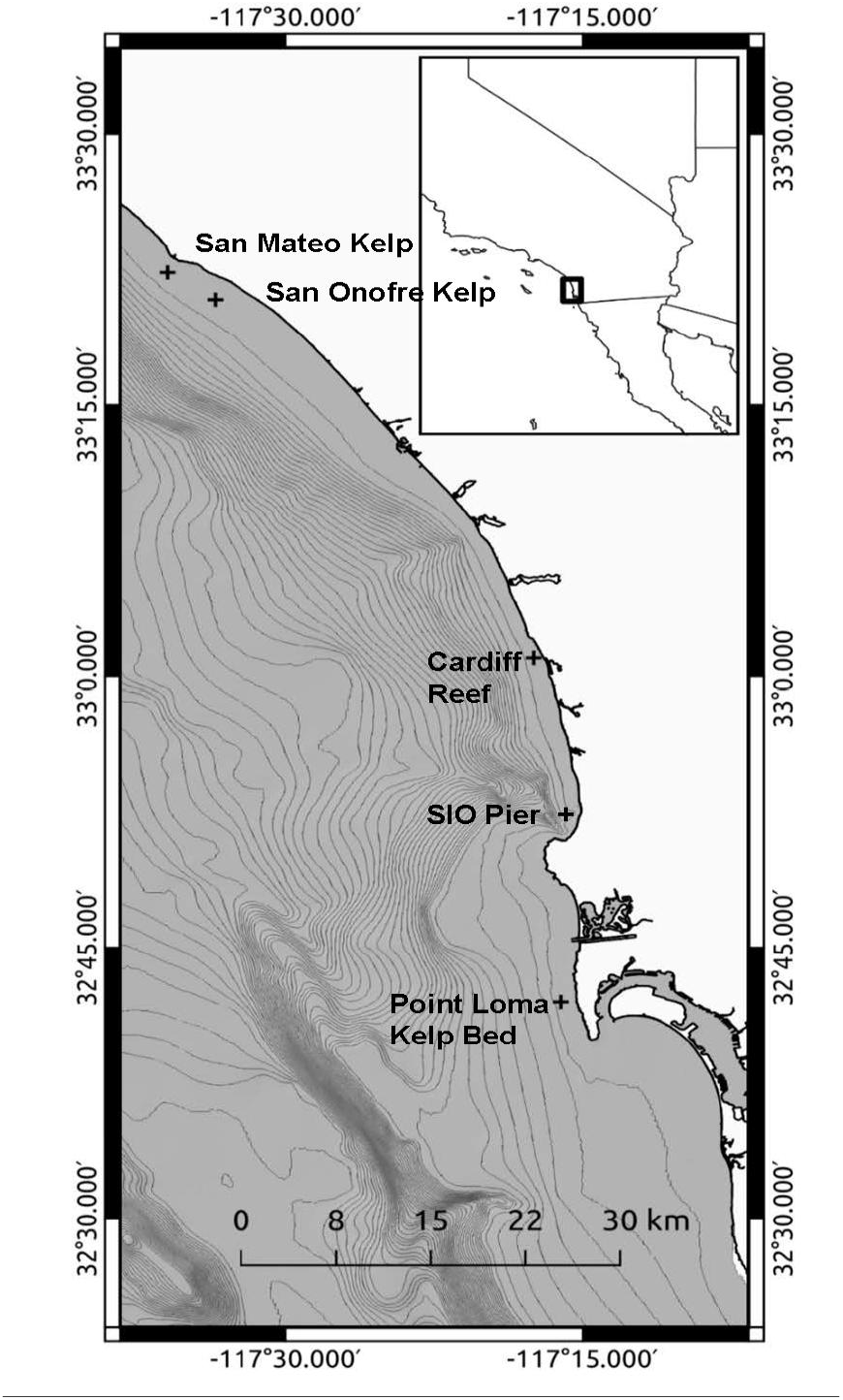
Map of study sites in southern Orange County, CA (San Mateo kelp Bed) and San Diego County, CA (San Onofre kelp bed, Cardiff Reef, Point Loma kelp bed, and Scripps Institution of Oceanograph (SIO) pier). Depth contour intervals are 30 meters.

### 2.2 Effects of Temperature

Two experiments were conducted to determine the effects of water temperature on symptoms of wasting disease in *Asterina miniata*. The first experiment used 18 *A. miniata* collected in the San Onofre kelp bed at a depth of 15 m on May 7, 1985. Sea stars were divided into two groups of nine. Size distributions were similar in the two groups and all 18 individuals were healthy-appearing and had no visible lesions. Nine of the sea stars were placed in aquaria where the temperature was maintained at 14°-17°C and nine were placed in aquaria where the temperature was maintained at 20°– 21°C. Individuals in both groups were observed daily from May 7 to May 22, 1985 for symptoms of disease.

The second experiment used 16 *A. miniata* collected on June 17, 1985, in the San Onofre kelp bed at a depth of 15 m. All individuals appeared healthy and had no visible lesions. The size distributions were similar in the two groups. Nine individuals were placed in aquaria where the temperature was maintained at 12°-13°C and seven were placed in aquaria where the temperature was maintained at 20°-21°C. Sea stars in both groups were observed daily from June 17 to June 27, 1985, for symptoms of disease.

### 2.3 Causative organism

An experiment was conducted to determine whether disease symptoms in *A. miniata* were associated with virus-sized or bacteria-sized organisms. Nineteen *A. miniata* and ambient seawater from the same area were collected at a depth of about 16 m in the offshore portion of the San Onofre kelp bed in late June 1985. These individuals were held in a seawater tank maintained at 18°C. Seven of the sea stars developed extensive lesions. Tissues from these diseased animals were homogenized in a blender and the homogenate was filtered through either a 0.5-micron pore-size Millipore filter, which allowed both bacteria-sized and virus-sized particles to pass through, or through a 0.2-micron pore-size filter, which allowed only virus-sized particles to pass through. On July 10, 1985, a challenge experiment was begun in which three groups of four of the 12 healthy-appearing *A. miniata* were injected with either the raw seawater, filtrate from the 0.5-micron filter, or filtrate from the 0.2-micron filter, examined daily for 14 days, and then scored as healthy, with wasting disease lesions, or dead.

In January 1987, we collected diseased *A. miniata* and *Henricia leviuscula* from the San Onofre kelp bed (SOK) and from the San Mateo kelp bed (SMK) in northern San Diego County (Figure 1). Eight bacterial isolations were cultured from the epidermal lesions of the diseased animals and identified using standard phenotypic methods (J.E. Hose, unpublished data). All were Vibrio species (one *V. harveyi*, two *V. nigripulcritudo*, one *V. alginolyticus*, one *Vibrio* sp. not further identified, and three *Vibrio* spp. (one from *H*. l*eviuscula* and two from *A. miniata*) with the same biochemical characteristics^1^ that were most similar to those of *V. metschnikovii* among described species in 1987. We will refer to these three isolates simply as *Vibrio* sp. A).

There were too few healthy-appearing *A. miniata* at the San Onofre and San Mateo kelp beds to use in a challenge experiment, Therefore, we searched other kelp beds along the San Diego County coast and on January 26, 1987 collected 31 healthy-appearing *A. miniata* from Bird Rock, a reef offshore of La Jolla, California. These animals were held in an aquarium maintained at 20°– 21°C for 16 days, during which no disease symptoms appeared. Individuals were then placed in separate aquaria and randomly assigned to treatment. Groups of three *A. miniata* were inoculated with each of the eight bacterial isolates obtained from diseased sea stars in the wild and seven were controls that were injected with seawater filtered through a 0.2-micron Millipore filter. The experimental animals were examined daily for two weeks and then scored as healthy, with wasting disease lesions, or dead.

On May 4, 1987, 28 healthy-appearing *A. miniata* were collected from Bird Rock and held at temperatures between 20° and 21° for 20 days, during which no disease symptoms appeared. Individuals were then placed in separate aquaria and randomly assigned to treatment. Fourteen *A. miniata* were inoculated with isolates of *Vibrio* sp. A obtained from the lesions of the sea stars that died of disease in the first challenge experiment and fourteen were controls that were injected with seawater filtered through a 0.2-micron Millipore filter. The experimental animals were examined daily for two weeks and then scored as either healthy, with wasting disease lesions, or dead. At the end of the experiment all 14 treated individuals were tested for *Vibrio* sp. A.

The last significant outbreak of wasting disease that we observed in the 1980s occurred in the late summer 1987 in a small patch of giant kelp in the San Onofre kelp bed at a depth of about 11 m. We collected *A. miniata* with necrotic lesions from which 17 bacterial isolations were cultured and identified using standard phenotypic methods described above (J.E. Hose, unpublished data).

### 2.4 Population sampling

From 1980 to 1987, as part of a study of the environmental effects of the San Onofre Nuclear Generating Station (Schroeter, et al. 1993), sea star density was estimated two to four times a year at two stations in the San Onofre kelp bed and at one station in the San Mateo kelp bed (Figure 1). At each sampling site, animals were counted in 40 1-m^2^ quadrats spaced 4 m apart along four 40-m transects arrayed in the shape of a cross with a common origin. From 2000 through 2020, as part of studies of an artificial reef constructed near the San Mateo kelp bed, sea star density was estimated annually within the San Mateo kelp bed (Reed, et al. 2023). Sea stars were counted in five 2 m x 10 m parallel plots equally spaced along each of 82 fixed 50 m x 20 m transects. The transects were arrayed in pairs 25 m apart uniformly placed throughout the kelp bed on rocky substrates known to have supported persistent giant kelp. The annual sampling was conducted from late spring to early fall.

Conspicuous invertebrates were censused annually during the spring in four permanently affixed 100-m^2^ band transects (25 m × 4 m) oriented perpendicular to shore at five sites at Point Loma and one site at Cardiff Reef. Each transect was composed of ten 2 m × 5 m quadrats within which invertebrates were counted. The means and standard errors for Point Loma are based on the combined data from all five sites.

## 3. RESULTS

### 3.1 Seawater temperature

We have included Scripps Institution of Oceanography data from the period 1980 through 2020 (Figure 2). Mean sea surface temperatures fluctuated over the course of our studies but showed increased temperature signals during El Niño periods in 1982-83, 1997-98, 2004-05 and 2014-16. During these periods of elevated temperatures, we also observed sea stars with wasting disease (Figure 2).

**Figure 2.**
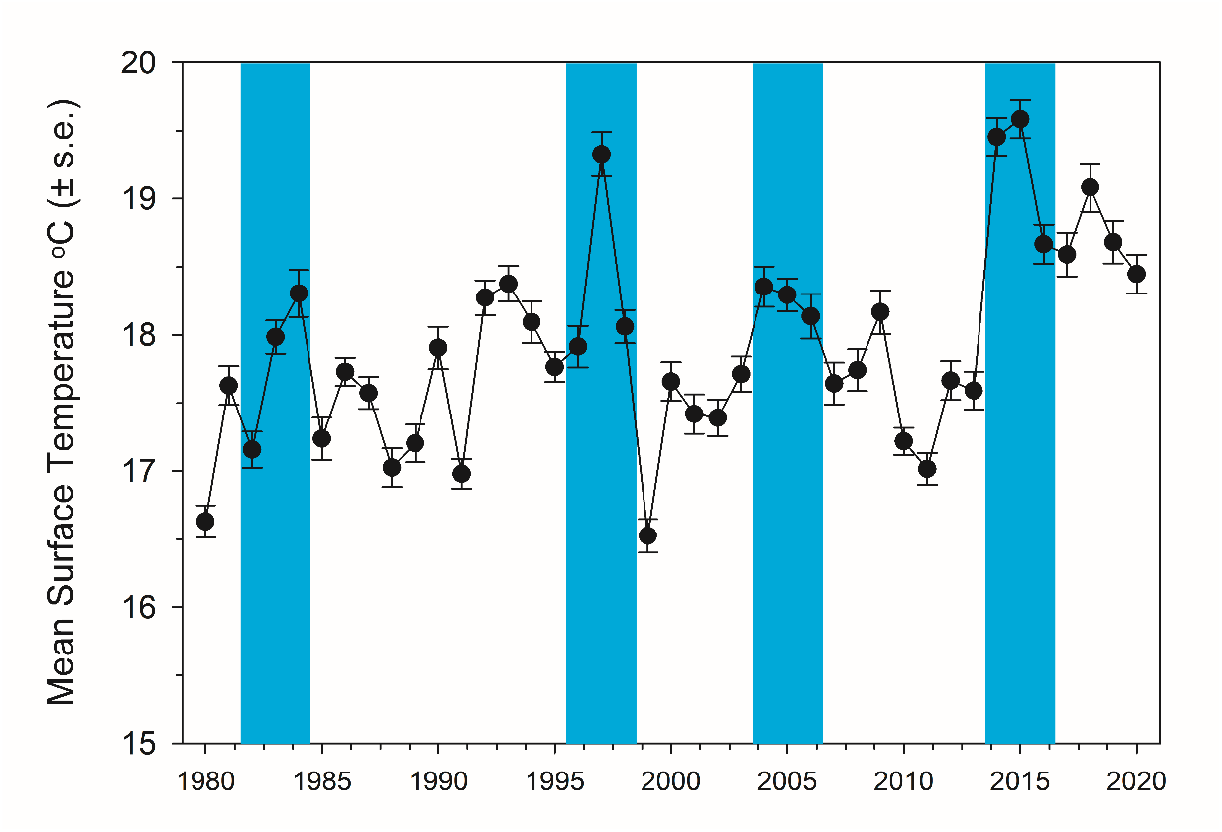
Average yearly sea surface temperatures at the Scripps Institution of Oceanography pier (Figure 1). Shaded blue bars indicate years during which diseased sea stars were observed and sea star wasting disease epizootics occurred. These periods included warm water events associated with El Niños.

### 3.2 Effects of temperature

Experiment 1 (May 7 – May 22, 1985): All nine of the sea stars held in aquaria at 14°-17°C remained healthy, whereas one of the nine individuals held at 20°-21°C developed disease symptoms. Experiment 2 (June 18 – June 27, 1985): Nine healthy-appearing *Asterina miniata* held in aquaria at 12°-13°C did not develop disease symptoms, whereas all seven of those held at 20°-21°C developed lesions and six died.

### 3.3 Causative Organism

Of the four *A. miniata* inoculated with the 0.2-micron filtrate, none developed disease symptoms, whereas three of the four inoculated with the 0.5-micron filtrate developed lesions typical of wasting disease and two died. Similarly, three of the four challenged animals inoculated with raw seawater developed lesions and one died, indicating that the infectious disease agent was water-borne.

In the challenge experiment, one experimental animal inoculated with *V. nigripulcritudo* and one inoculated with *V. alginolyticus* developed small lesions, but neither died. None of the other isolates resulted in disease symptoms except for the three *Vibrio* sp. A isolates. *Vibrio* sp. A produced typical lesions in four of the nine challenged animals and three of these died. Nine healthy-appearing *A. miniata* injected with seawater filtered through a 0.2-micron filter developed no disease symptoms. Of the fourteen *A. miniata* that were inoculated with isolates of *Vibrio* sp. A obtained from the lesions of the sea stars that died of disease in the first challenge experiment, four developed lesions and died. All 14 of the control animals remained healthy.

By 1987 there were very few sea stars remaining at our study sites. Of the 17 bacterial isolations developed from diseased *A. miniata* collected in the San Onofre kelp bed in late summer 1987, 13 or 77% were *Vibrio* sp. A.

These experimental and sampling data strongly indicate that *V*ibrio. sp. A causes wasting disease in *Asterina miniata* and *Henricia leviuscula* and was the causative agent in the epizootics that affected those species, and probably all the affected sea stars, in the 1980s. Based on the species affected and the very similar disease symptoms and coarse of the epizootics during the El Niños in 1997 and 2004, it is likely that Vibrio sp. A was the causative agent then too. The fact that many of the challenged individuals did not develop disease symptoms was surprising. However, those experiments were conducted toward the end of the epizootic when there were relatively few surviving sea stars. We speculate that the survivors tended to have some immunity to the disease.

### 3.4 Population Changes

During the period 1980 to 1987, at least seven species of sea stars developed the wasting disease at our San Mateo and San Onofre study sites: *Astropecten armatus, Linckia columbiae, Dermasterias imbricata, Henricia leviuscula, Asterina miniata, Astrometis sertulifera*, and *Pisaster giganteus. Asterina miniata* and *Pisaster giganteus* were most abundant. Observations at the Point Loma kelp bed were similar, with the exception of *Astropecten armatus* and *Linckia columbiae* that did not occur there. These observations of wasting disease generally occurred during the periods of higher seawater temperatures associated with the El Nino events of 1982-83, 1997-98, 2004-05 and 2014-16 (Figure 2). During these warm water episodes, we also observed similar symptoms in the common holothurian *Parastichopus parvimensis* and in the echinoids *Strongylocentrotus franciscanus, S. purpuratus, Lytechinus anamesus*, and *Centrostephanus coronatus*, either on the mainland coast or at Catalina Island. From 2013 to 2015, the sea stars *Asterina miniata, Pisaster giganteus*, and the much less abundant *Henricia leviuscula* were observed with wasting disease at our study sites.

We have plotted the densities of *Asterina miniata* in the San Onofre and San Mateo kelp beds from 1980-1987, in the San Mateo kelp bed from 2000-2020, and in the Point Loma kelp bed from 1980-2020 (Figure 3). The most dramatic effect of the wasting disease was seen at the San Onofre kelp bed where *A. miniata* declined from >100 100 m^-2^ in 1981 to near zero in 1984. There was also a decline in density in the San Mateo kelp bed where *A. miniata* dropped from around 14 100 m^-2^ in 1981 to near zero in 1984. Although there was also a modest decline in numbers in the San Mateo kelp bed from 2003-2005 that may have been due to disease, the most striking pattern after 2000 was a buildup of numbers at both San Mateo and Point Loma followed by a precipitous decline during the 2014-2015 El Niño. There was some recovery at San Mateo, but not at Point Loma.

**Figure 3.**
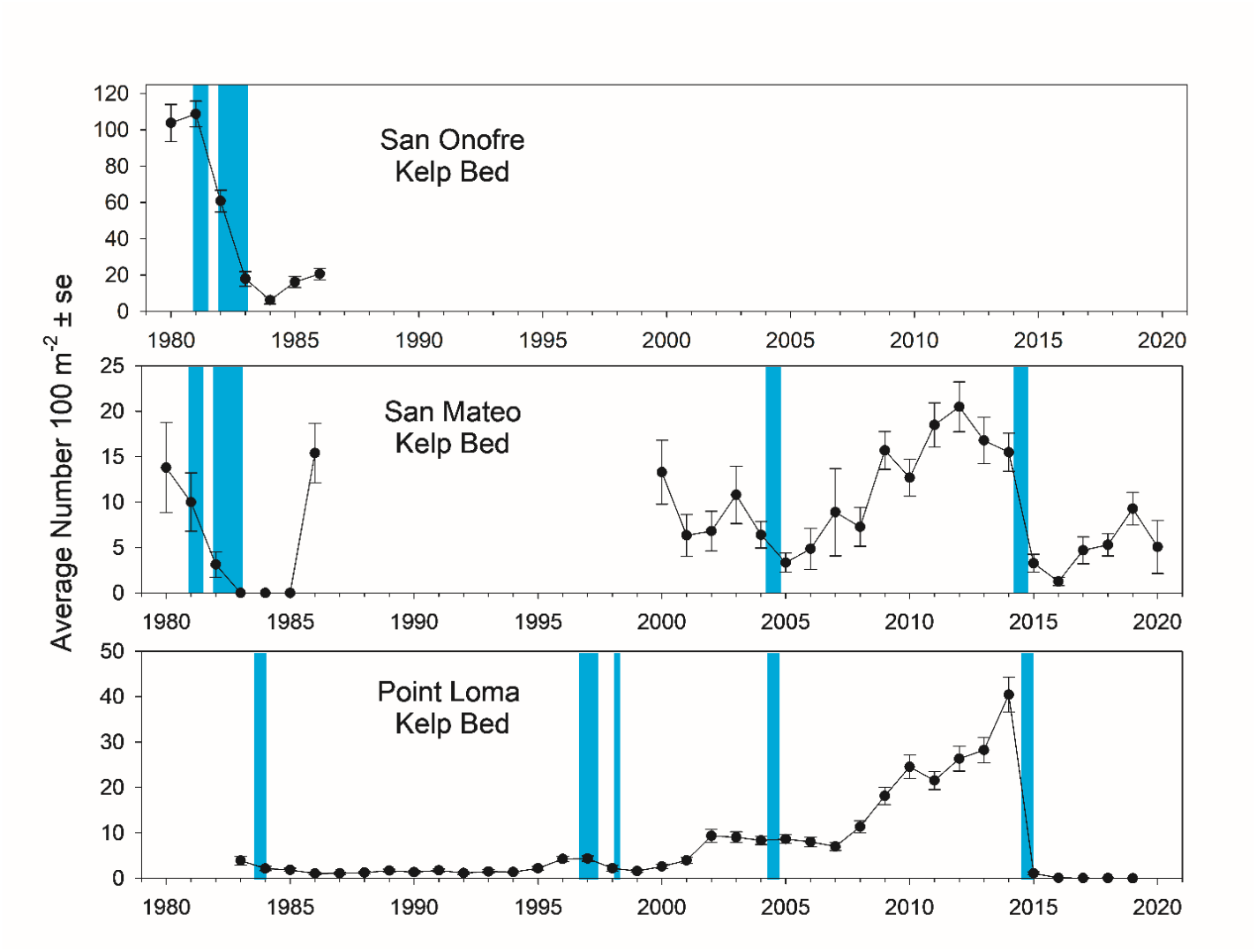
Mean density (+/- 1 s.e.) of *Asterina miniata* in the San Onofre, San Mateo, and Point Loma kelp beds from 1980 to 2020. Colored vertical bars indicate periods when wasting disease was observed in the field.

Similar patterns of decline in abundance concomitant with observations of wasting disease were observed for *P. giganteus* in the San Mateo and Point Loma kelp beds where densities remained above ten 100 m^-2^ until 1983 and then declined to less than five 100 m^-2^ in 1985. From 1985-1997 at Point Loma there was recovery followed by decline to near zero during the 1997-1998 El Niño. From 2000 until 2014 abundances fluctuated a great deal at San Mateo, Point Loma, and Cardiff Reef but increased from 2006 to 2014 followed by a sharp drop to zero after 2014 (Figure 4).

**Figure 4.**
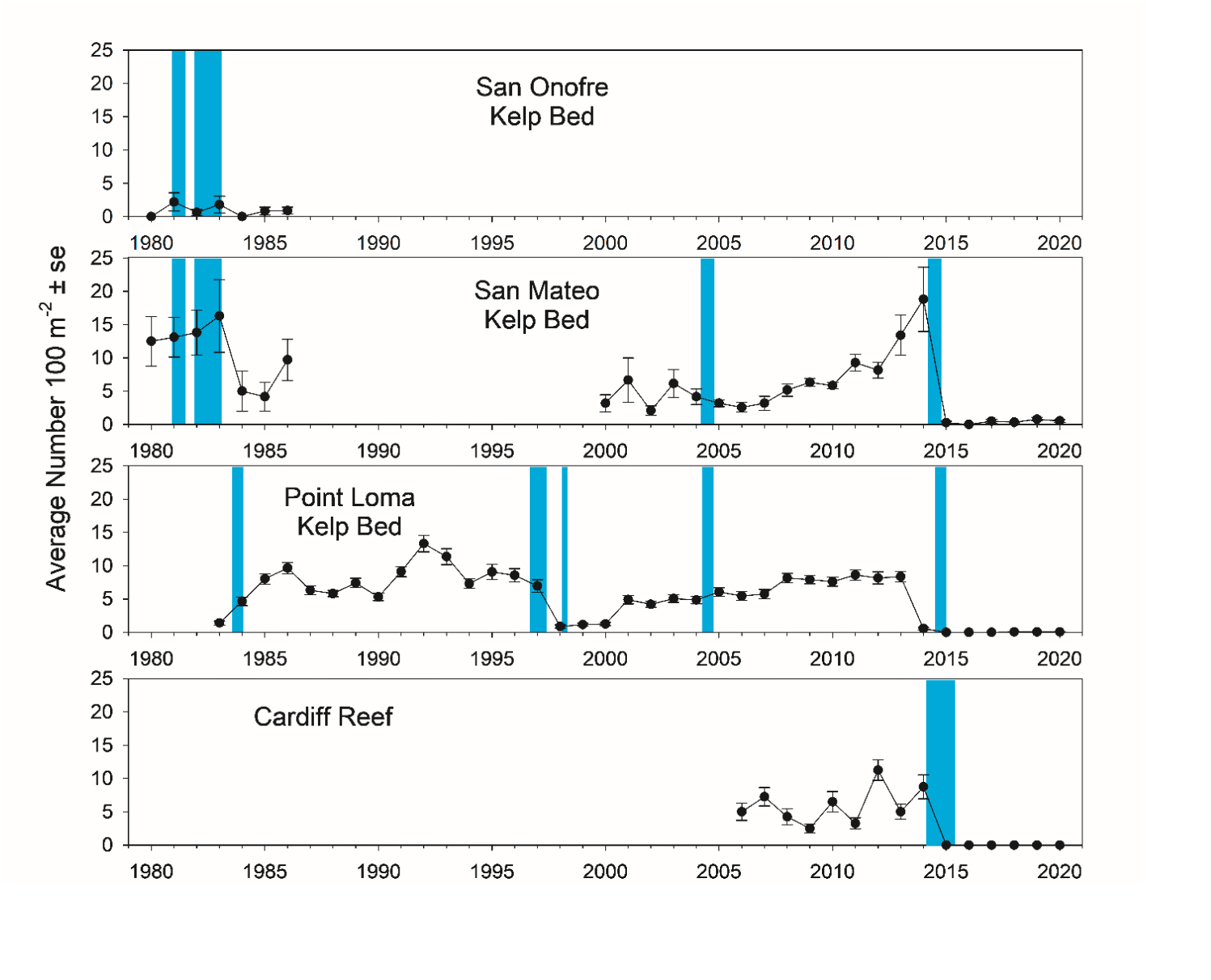
Mean density (+/- 1 s.e.) of *Pisaster giganteus* in the San Onofre, San Mateo, Point Loma, and Cardiff Reef kelp beds. Colored vertical bars indicate periods when wasting disease was observed in the field.

Densities of *Pisaster brevispinus* in the Point Loma kelp bed exhibited a similar pattern through time. There were declines during warm water periods followed by recovery prior to the marine heat wave beginning in 2014 when there was a sharp decline to zero. Since 2015, there has been essentially no recovery of any of the species we have monitored (Figures 3-5). The one exception is *Asterina miniata* in the San Mateo kelp bed where densities plummeted during the 2014-2016 El Niño, but have since fluctuated from around 1 to 10 100 m^-2^.

**Figure 5.**
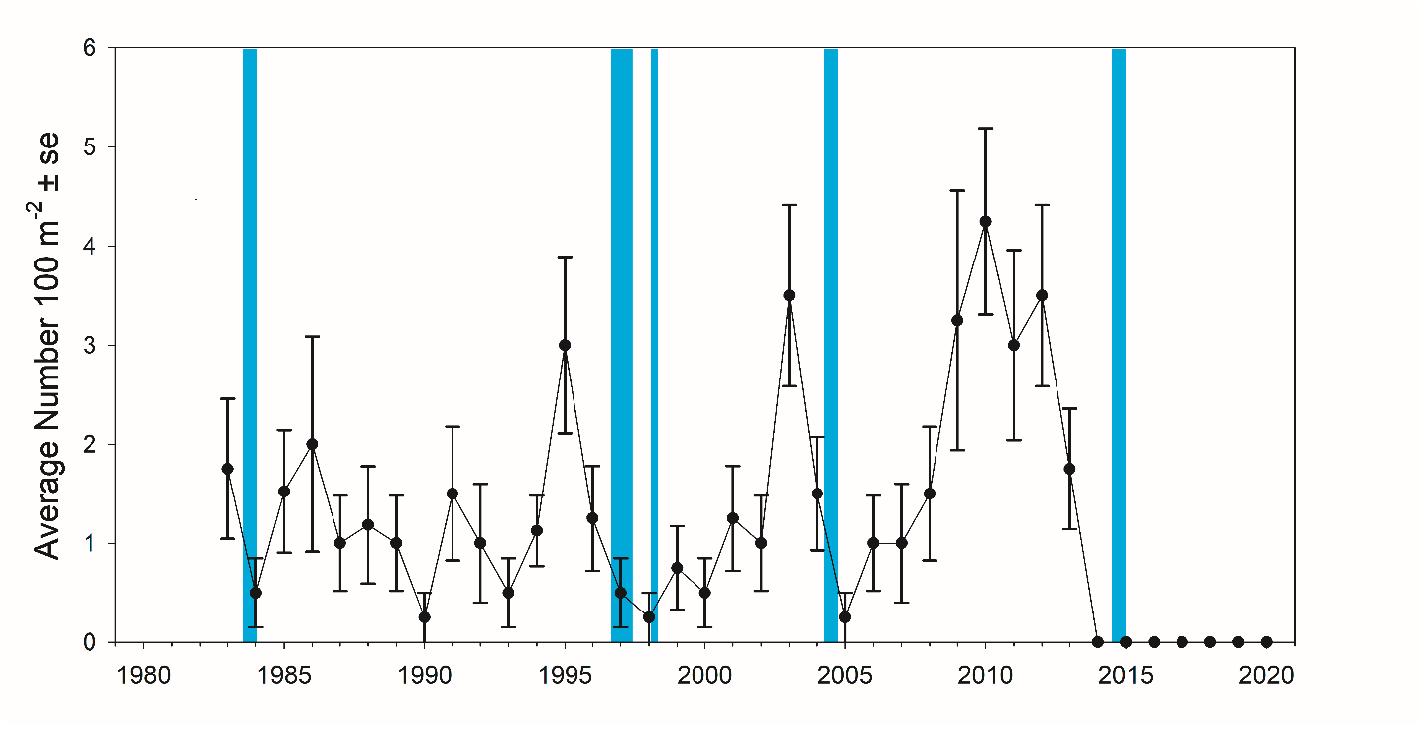
Mean density (+/- 1 s.e.) of *Pisaster brevispinus*, in the Point Loma kelp bed. Colored vertical bars indicate periods when wasting disease was observed in the field.

## 4. DISCUSSION

Most of the sea stars present at our study sites during the 1981-1984 epizootic developed lesions and these individuals completely decomposed and died within a few days. The resultant decline in numbers was precipitous. From 1981 to 1984, the population of *Asterina miniata* in the San Onofre kelp bed declined from an average of about 130 100 m^-2^ to about 3 100 m^-2^ as a result of at least three disease episodes (Figure 3). By 1987, the density of sea stars at all our study sites in northern San Diego County was close to zero. A similar pattern occurred in the San Mateo and Point Loma kelp beds for *P. giganteus* early in the time series. Densities were low in 1983 followed by an increase from 1983 to 1987, slow recovery through 2014, followed by a sharp drop after 2014. We observed a similar pattern at the Cardiff Reef kelp bed, namely variable levels of abundance until 2014, followed by a sharp drop. In all cases, these declines in abundance coincided with high sea water temperatures and observations of disease (i.e. lesions or decomposing sea stars).

The mass mortality of *Asterina miniata* resulted in a trophic cascade affecting both grazers and algae. Predation by *A. miniata* appears to control the distribution of the white sea urchin, *Lytechinus anamesus* (Schroeter et al. 1983). At our study sites, *A. miniata* usually occurred at higher densities inside the kelp beds where its preferred boulder substrate is more abundant than farther offshore. As a result, *L. anamesus* was only present in large aggregations along the offshore margins of the beds where they grazed on early developmental stages of kelps, preventing the offshore expansion of the kelp bed (Dean et al. 1984). Following the mass mortality of *A. miniata*, the numbers of *L. anamesus* increased substantially within the kelp beds where they altered their feeding behavior and began grazing on adult giant kelp (*Macrocystis pyrifera*), resulting in significant mortality. A similar trophic cascade was observed in Howe Sound, British Columbia where the sunflower star, *Pycnopodia helianthoides*, an abundant predator of subtidal bottom-dwelling invertebrates, suffered a nearly 90% decline in abundance from wasting disease. Following this mass mortality, green sea urchin abundance quadrupled and kelp (*Agarum finbriatum*) declined significantly (Schultz, et al., 2016).

Changes in community composition as a result of changes in the density of higher trophic level species are probably common (Estes, et al. 2011) and have been demonstrated experimentally (e.g., Paine 1966, 1974; Dayton 1971) and documented when natural causes reduce predator or herbivore density (e.g., Pearse and Hines 1979, Lessios 1988, Estes and Palmisano 1974, and Vicknair and Estes 2012).

Mass mortality events that devastate populations of marine organisms, often on a large spatial scale, appear to be increasing throughout the world (Harvell et.al 1999, Harvell 2019, Ward and Lafferty 2004). However, the causative agents are often not known. Our results indicate that a bacteria-sized microorganism, identified as a species of *Vibrio*, was the causative agent of the wasting disease epizootic that caused mass mortality at our study sites in the 1980s.

The 40-year time period of our study provides a unique opportunity to observe a number of marine heat waves and to compare and contrast their effects on the target sea star species. Unlike the events prior to the 2014-2016 heat wave, there was no recovery over the 6-year period following the dramatic declines in 2014, suggesting the possibility of increasing severity and persistence of disease on sea star populations may be a consequence of the wider geographical extent of the 2014 disease episode. The best estimate of planktonic larval dispersal times of the sea stars in Southern California is about 90 days (Reed et. al. 2000), likely enabling sea star populations to recolonize from nearby sites during the earlier, more geographically restricted epizootics. We speculate that the lack of recovery we’ve seen following the 2014-2016 marine heat wave may be the consequence of significantly reduced larval sources for recolonization following these geographically widespread epizootics.

## 6. ACKNOWLEDGEMENTS

We are particularly grateful to J. H. Hose for the bacterialogical work in the 1980s and for reviewing the manuscript. Partial funding for the 1980’s research was provided by faculty research grants to S. Schroeter and J. Dixon from the Division of Natural Sciences and Mathematics, College of Letters, Arts and Sciences, University of Southern California, and from the Marine Review Committee, Inc. (MRC), the entity overseeing research on the ecological effects of the San Onofre Nuclear Generating Station. Funding for the 2000’s research was provided by the San Onofre Nuclear Generating Station (SONGS) Mitigation Monitoring Program, Marine Science Institute, University of California Santa Barbara. Funding for the Pt. Loma kelp bed observations was provided by the National Science Foundation, and the City of San Diego.

## Footnotes

1 Yellow colonies on TCBS (thiosulfate citrate bile salt) agar, motile, halophilic facultative anaerobes utilizing sucrose (67% positive), uniformly negative for oxidase reaction and for gas production during glucose fermentation, sensitive to both 10 μg and 150 μg of vibriostatic agent O/129, and positive for B-galactosidase activity with ONPG (O-nitrophenyl-β-D-galactopyranoside) (J.E. Hose, unpublished data)

